# EEG data quality in large scale field studies in India and Tanzania

**DOI:** 10.1101/2024.12.05.626951

**Authors:** John-Mary Vianney, Shailender Swaminathan, Jennifer Jane Newson, Dhanya Parameshwaran, Narayan Puthanmadam Subramaniyam, Swaeta Singha Roy, Revocatus Machunda, Achiwa Sapuli, Santanu Pramanik, John Victor Arun Kumar, Pramod Tiwari, G Nelson Mathews Mathuram, Laurent Boniface Bembeleza, Joyce Philemon Laiser, Winifrida Julius Luhwago, Theresia Pastory Maduka, John Olais Mollel, Neema Gadiely Mollel, Adella Aloys Mugizi, Isaac Lwaga Mwamakula, Raymond Edwin Rweyemamu, Upendo Firimini Samweli, James Isaac Simpito, Kelvin Ewald Shirima, Anand Anbalagan, Suresh Kumar Arumugam, Vinitha Dhanapal, Kanimozhi Gunasekaran, Neelu Kashyap, Dheeraj Kumar, Durgesh Pandey, Poonam Pandey, ArunKumar Panneerselvam, Sonam Rai, Porselvi Rajendran, Santhoshkumar Sekar, Oliazhagan Sivalingam, Prahalad Soni, Pushpkala Soni, Tara C. Thiagarajan

## Abstract

There is a growing imperative to understand the neurophysiological impact of our rapidly changing and diverse technological, social, chemical, and physical environments. To untangle the multidimensional and interacting effects requires data at scale across diverse populations, taking measurement out of a controlled lab environment and into the field. Electroencephalography (EEG), which has correlates with various environmental factors as well as cognitive and mental health outcomes, has the advantage of both portability and cost-effectiveness for this purpose. However, with numerous field researchers spread across diverse locations, data quality issues and researcher idle time due to insufficient participants can quickly become unmanageable and expensive problems. In programs we have established in India and Tanzania, we demonstrate that with appropriate training, structured teams, and daily automated analysis and feedback on data quality, non-specialists can reliably collect EEG data alongside various survey and assessments with consistently high throughput and quality. Over a 30-week period, research teams were able to maintain an average of 25.6 subjects per week, collecting data from a diverse sample of 7,933 participants ranging from Hadzabe hunter-gatherers to office workers. Furthermore, data quality, computed on the first 2,400 records using two common methods, PREP and FASTER, was comparable to benchmark datasets from controlled lab conditions. Altogether this resulted in a cost per subject of under $50, a fraction of the cost typical of such data collection, opening up the possibility for large-scale programs particularly in low- and middle-income countries.

**Significance Statement:** With wide human diversity, a rapidly changing environment and growing rates of neurological and mental health disorders, there is an imperative for large scale neuroimaging studies across diverse populations that can deliver high quality data and be affordably sustained. Here we demonstrate, across two large-scale field data acquisition programs operating in India and Tanzania, that with appropriate systems it is possible to generate high throughput EEG data of quality comparable to controlled lab settings. With effective costs of under $50 per subject, this opens new possibilities for low- and middle-income countries to implement large-scale programs, and to do so at scales that previously could not be considered.

## Introduction

The past 50 years have witnessed an accelerated transformation of our technological, social, cultural and physical environment (Arora, 2019, Anon, 2019; Roser et al., 2024). However, how these changes impact our brain physiology is still poorly understood. As an experience-dependent organ, the human brain is sensitive to change and variation in our stimulus environment. For example, electroencephalography (EEG) studies have demonstrated differences in resting-state and evoked potentials in response to inter-individual differences in demographic profiles (Tomescu et al., 2018; Sandre et al., 2024), lifestyle habits (Khoo et al., 2024), developmental stages (Anderson and Perone, 2018; Wilkinson et al., 2024), and stimulus (Parameshwaran et al., 2019, 2021; Parameshwaran and Thiagarajan, 2023) or physical environments (Hou et al., 2023).

Understanding of the changing impacts of modern life on the brain requires large-scale high throughput studies that acquire data across diverse cross-sections of a population. While there are several larger-scale studies in progress, such as the Adolescent Brain Cognitive Development Adolescent (ABCD^®^) Study (Casey et al., 2018), Human Connectome Project (Van Essen et al., 2013), UK Biobank (Miller et al., 2016), Cuban Human Brain Mapping Project, CHBMP (Valdes-Sosa et al., 2021), Child Mind Institute’s’ Healthy Brain Network (Alexander et al., 2017) and ENIGMA Consortium (Thompson et al., 2020), they are typically resource intensive, or require fixed infrastructure, and consequently are able to acquire samples only on the scale of 10,000 or less, are geographically limited and therefore and do not reflect the breadth of human environment or culture. The ABCD project (Casey et al., 2018), for example, which utilizes fMRI as its primary neuroimaging device and captures data across 11,000 children each year in the United States has an annual budget of $41 million, which is prohibitive for most low- and middle-income countries. While affordable electroencephalography (EEG) devices are now available on the order of a few thousand dollars, a major aspect of cost is the need for trained or specialized technicians and research scientists, as well as ensuring sufficient throughput that minimizes idle time.

The only way such large-scale programs can be established affordably are if non-specialists can be entrusted to collect data with a high throughput of participants coupled with high data quality. With many field researchers working across various field locations, data quality issues with respect to both surveys and physiological recordings can quickly become an unmanageable problem. This can be overcome by establishing robust systems and processes. First, effective recruitment and training of field research staff, second, daily dashboards and feedback on data quality to identify challenges as soon as they arise rather than after they have perpetuated, and third, effective mechanisms for recruitment of participants and coordination of logistics in the field.

Sapien Labs has established and piloted such scalable systems and processes through its Centers for Human Brain and Mind at Krea University in India and the Nelson Mandela African Institute of Science and Technology (NM-AIST) in Tanzania where non specialist field researchers capture survey, task and EEG data across diverse field locations using a low-cost EEG device. Here we describe the data throughput and resulting EEG data quality achieved in these programs, from the first 1,100 and 1,300 participants across India and Tanzania, respectively. EEG data quality challenges primarily include eye blink and movement artifacts and power line noise, but can also include fluctuations in impedance and challenges with electrode placement due to varying hairstyles. We used two commonly available pipelines (Preprocessing Pipeline (PREP) (Bigdely-Shamlo et al., 2015) and Fully Automated Statistical Thresholding (FASTER) (Nolan et al., 2010), to measure the percentage of bad channels and bad epochs, and compared the results against EEG data from 3 highly cited benchmark datasets with equivalent experimental tasks obtained in a controlled lab environment with more expensive EEG devices (Singh et al., 2022; Wang et al., 2022; Miltiadous et al., 2023; Anjum et al., 2024; Xiang et al., 2024).

## Materials and Methods

### EEG equipment

EEG was recorded with the wireless Emotiv FLEX 2 Gel headset using 16 out of 32 electrodes positioned according to the 10–20 International system and referenced to an ear clip sensor. The montage included 8 electrodes over each hemisphere with alternative configurations utilized on occasion when hair type or style imposed a restriction of data acquisition (e.g. braided buns) (Figure 1). The internal sampling rate was 2048 samples per second downsampled to 256 Hz.

**Figure 1:**
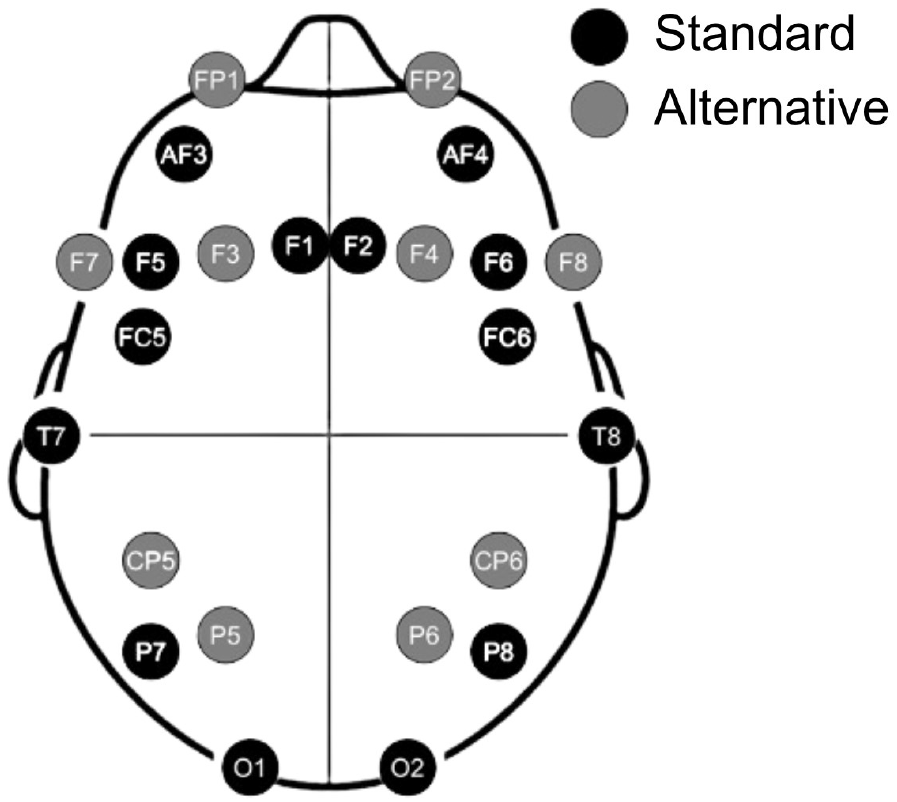
Electrode configuration used in the project. Black = standard configuration; grey = alternate non-standard channels allowed in case of hair obstruction of a channel.

### Research Personnel

Field researchers were recruited among new college graduates as well as people with experience in field survey methods. All researchers were EEG naïve and were trained in the use of the EEG device for two weeks prior to field data collection. A total of 12 field researchers were recruited in each country (India and Tanzania) and trained over two weeks. On day 1, trainees were given a demonstration and explanation of EEG and performed hands on trials with the help of a trainer who was experienced in EEG data acquisition. On days 2 and 3, trainees performed hands-on resting-state EEG trials in pairs without direct assistance of the trainer and participated in problem solving and debriefing sessions throughout the day. Each trainee collected EEG data from 4-6 people over the two days and received immediate feedback on data quality. Trainees then spent one week in the field, recording EEG from 10-15 participants under various circumstances (open air rural locations, office rooms etc.) and undertook debrief sessions at the end of the day to discuss challenges relating to field settings, equipment and data quality, and identify solutions. Throughout the training, trainees were evaluated for their proficiency by the trainer. Trainees who were unable to reach minimum data quality and throughput numbers by the end of the training period were not hired. After the training, 93% met the standard and were retained.

### Team structure

The 12 field researchers in each country were divided into teams of 2 or 3. In addition, one participant recruitment manager (PRM) worked with field researchers in each region/country. The PRMs were recruited based on strong local networks as well as communication and organization skills and were responsible for identifying study participants and locations to fit with the sampling frame, reaching out to participants and ensuring that they were able to report to the study location at a specified time, as well as ensuring local government permissions and all logistics.

### Participant recruitment

Study participants were recruited in multiple locations in India and Tanzania according to a sampling frame designed to cover a broad range of income groups (low, medium and high) across the lifespan, divided equally across biological males and females and divided among different types of geographies and settlements. In India, this covered multiple regions in the southern state of Tamil Nadu as well as the National Capital Region (NCR; Delhi, Haryana, Rajasthan, and Uttar Pradesh). In Tanzania, target locations spanned Arusha and Manyara regions including rural, suburban and urban areas. Participants were age 18+ in Tanzania and age 13+ in India. Recordings were carried out in various locations including offices, schools, and open air.

All participants gave written informed consent and all procedures involving human subjects were approved by an ethical review board [India: IFMR Institutional Ethics Committee (IEC) IFMR-IHEC/SL/0001/2023; National Ethics Committee Registry for Biomedical and Health Research (NECRBHR), Department of Health Research (DHR); EC/NEW/INST/2023/3887; Tanzania: Kibong’oto Infectious Diseases hospital - Nelson Mandela African Institution of Science and Technology – Centre for Educational Development in health, Arusha (KIDH-NM-AIST-CEDHA) – KNCHREC; KNCHREC00006/09/2023].

### Experimental design and statistical analysis

#### EEG protocol

Resting-state EEG was collected when participants were sitting quietly with their eyes closed (EC) for 3 minutes and eyes open (EO) for 3 minutes. During the EO task, participants were instructed to look at their surroundings, rather than the laptop or the researcher. In addition, all participants completed a Raven’s progressive matrix task (Raven, 2000). Participants completed various surveys including a session information questionnaire where they reported any physical symptoms (e.g. headache, cold, stomach ache), any medications they had taken in the past 24 hours, any substances such as caffeine or drugs consumed in the past 12 hours, time of last meal, duration of previous night’s sleep and time since they woke up as well as their mood and alertness at the time of recording.

#### Data quality monitoring

Data quality was monitored in real time by researchers with end of day reports returned to each field researcher. In addition, the channel quality metrics available in Emotiv’s recording software, scripts were run on test data obtained after electrode positioning to compute data quality and indicate any adjustments needed. Experimental protocols were initiated only after the test signal passed this quality test. In addition, post recording, all data were automatically analyzed for signal quality using the FASTER Z-score criteria. A dashboard showing both throughput and signal quality could be viewed by the field researchers and supervising staff for immediate course correction in case of arising data quality issues (Table 1).

**Table 1:**
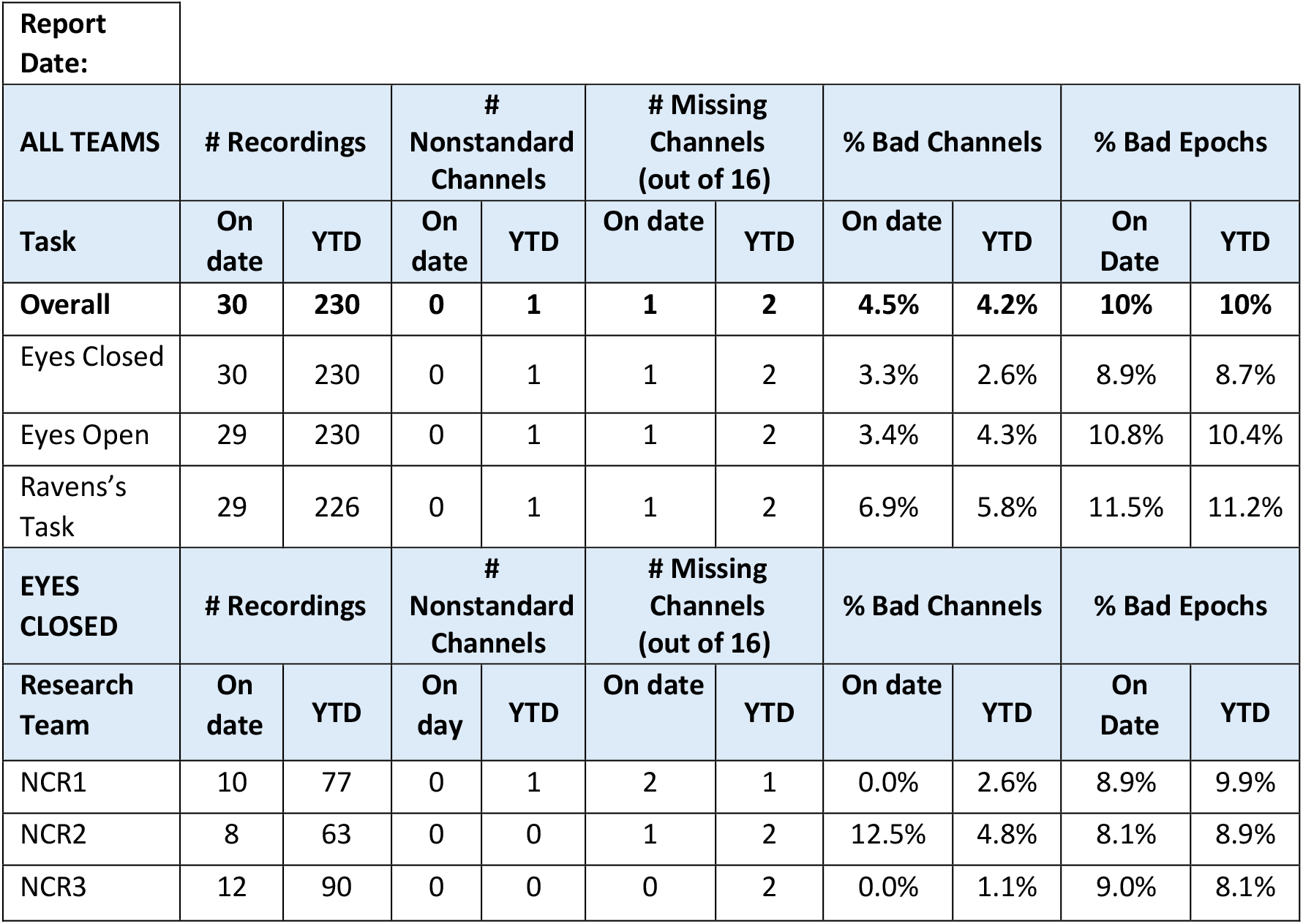
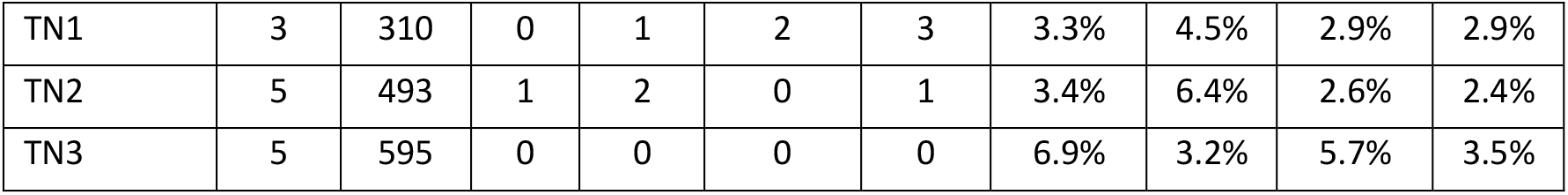
Daily report provided to field research team and supervisors in India. TN: Tamil Nadu Team; NRC: National Capital Region; YTD: Year to Date.

| Report Date: |  |  |  |  |  |  |  |  |  |  |
| --- | --- | --- | --- | --- | --- | --- | --- | --- | --- | --- |
| ALL TEAMS | # Recordings |  | # Nonstandard Channels |  | # Missing Channels (out of 16) |  | % Bad Channels |  | % Bad Epochs |  |
| Task | On date | YTD | On date | YTD | On date | YTD | On date | YTD | On Date | YTD |
| Overall | 30 | 230 | 0 | 1 | 1 | 2 | 4.5% | 4.2% | 10% | 10% |
| Eyes Closed | 30 | 230 | 0 | 1 | 1 | 2 | 3.3% | 2.6% | 8.9% | 8.7% |
| Eyes Open | 29 | 230 | 0 | 1 | 1 | 2 | 3.4% | 4.3% | 10.8% | 10.4% |
| Ravens's Task | 29 | 226 | 0 | 1 | 1 | 2 | 6.9% | 5.8% | 11.5% | 11.2% |
| EYES CLOSED | # Recordings |  | # Nonstandard Channels |  | # Missing Channels (out of 16) |  | % Bad Channels |  | % Bad Epochs |  |
| Research Team | On date | YTD | On day | YTD | On date | YTD | On date | YTD | On Date | YTD |
| NCR1 | 10 | 77 | 0 | 1 | 2 | 1 | 0.0% | 2.6% | 8.9% | 9.9% |
| NCR2 | 8 | 63 | 0 | 0 | 1 | 2 | 12.5% | 4.8% | 8.1% | 8.9% |
| NCR3 | 12 | 90 | 0 | 0 | 0 | 2 | 0.0% | 1.1% | 9.0% | 8.1% |
| TN1 | 3 | 310 | 0 | 1 | 2 | 3 | 3.3% | 4.5% | 2.9% | 2.9% |
| TN2 | 5 | 493 | 1 | 2 | 0 | 1 | 3.4% | 6.4% | 2.6% | 2.4% |
| TN3 | 5 | 595 | 0 | 0 | 0 | 0 | 6.9% | 3.2% | 5.7% | 3.5% |

**Table 2:** Details on benchmark datasets BM1, BM2 and BM3 used in this study to compare EEG data quality.

| Condition | BM1 | BM2 | BM3 |
| --- | --- | --- | --- |
| Eyes Open | Three minutes of 64channel EEG from 49 participants (Anjum et al., 2024) | Five minutes of 64channel EEG data from 60 participants (Wang et al., 2022) | Five minutes of 61-channel EEG data from 71 participants (Xiang et al., 2024) |
| Eyes Closed | 19-channel EEG recordings from 29 participants (Miltiadous et al., 2023) | Five minutes of 64channel EEG data from 60 participants (Wang et al., 2022) | Five minutes of 61-channel EEG data from 71 participants (Xiang et al., 2024) |
| Task | 64-channel EEG data from 60 subjects performing mathematics task (subtraction) (Wang et al., 2022) | 64-channel EEG data from 60 subjects performing mathematics task (Memory task involving recollecting events of the day) (Wang et al., 2022) | 69-channel EEG from 23 subjects performing working memory task involving memorizing and ignoring a set of presented letters (modified Sternberg task) (Singh et al., 2022) |

#### EEG data quality analysis

The quality of EEG recordings from India (EC: N=1,349; EO: N=1,341; Task: N = 1,329) and Tanzania (EC: N=1,168; EO: N=1,167; Task: N = 1,142) were evaluated using two commonly accepted approaches: (1) FASTER (Nolan et al., 2010) and (2) PREP (Bigdely-Shamlo et al., 2015). Each EEG recording was evaluated for percentage of bad epochs and bad channels based on the criteria proposed by the FASTER and PREP methods. For detecting bad epochs, the EEG data were divided into epochs of 2 seconds. The EEG recordings were high-pass filtered at 0.5 Hz before the detection of bad channels and epochs.

##### Detection of bad channels by FASTER

Detection of bad channels was based on three parameters (Nolan et al., 2010) which included:

1. A mean correlation coefficient between channels pairs with Z-score > 3 implying non-EEG signal contamination.
2. A signal variance Z-score > 3.
3. Hurst-exponent with Z-score > 3.

In addition to these three criteria, we also assessed contamination with powerline noise. To this end, channels with a mean power Z-score > 3 between 48-62 Hz were flagged as bad channels.

##### Detection of bad channels by PREP

Criteria used in the PREP method to identify bad or unusable channels was based on:

1. EEG channels with flat signals (threshold < (< 1e-15 *μV*) and NaN values, or channels with a flat-signal (< 1e-15 *μV*) for more than 1% of windows.
2. Amplitudes that exceeded a robust Z-score > 5 [as compared to standard Z-score by FASTER method (Nolan et al., 2010).]
3. A correlation threshold of < 0.4 between more than 1% of all 2-second windows in the signal.
4. The ratio of high (>50 Hz) and low frequency components exceeding a robust Z-score of 5 where a 50 Hz lowpass finite impulse response (FIR) filter was used to separate the low and high frequency components.

##### Detection of bad epochs by FASTER

Criteria used by the FASTER method was based on:

1. An amplitude range transformed Z-score > 3, where amplitude range was calculated as the difference between the maximum and minimum value in each epoch.
2. Variance within an epoch having a Z-score > 3 (used in order to detect artifacts due to participant movement).
3. Z-score of the deviation parameter for an epoch > 3, where deviation parameter measured the deviation of an epoch’s average value (across time) from the average values across all channels. For N epochs, this resulted in N x M deviation values, where M was the number of EEG channels. The deviation parameter values were then averaged across M channels resulting in N deviation parameters.

##### Detection of bad epochs by PREP

We modified the PREP method (Bigdely-Shamlo et al., 2015) to also detect bad epochs. For each EEG channel and epoch, robust standard deviation was computed using the interquartile range and multiplying it by 0.7413. For each channel and epoch, the median values were also computed, following which a robust Z-score was obtained for each epoch in each channel. A maximum robust Z-score > 5 for each epoch across all the channels was marked as a bad epoch.

### Benchmark EEG datasets

To facilitate the comparison of the quality of our EEG recordings with EEG datasets acquired in a more controlled setting, we compared the results of EEG quality metrics for each condition (EO, EC or TASK) against benchmark datasets obtained from OpenNeuro or NEMAR which we refer to as BM1, BM2 and BM3 (Singh et al., 2022; Wang et al., 2022; Miltiadous et al., 2023; Anjum et al., 2024; Xiang et al., 2024). These represent highly cited datasets that are openly available and described in Table 1.

## Results

Figure 2 shows the weekly throughput per EEG device for the first 30 weeks of data collection for the India and Tanzania teams. Weekly throughput was calculated as the number of participants recorded per EEG device per week since the first week post-training, where each device was managed by a team of 2-3 field researchers (average 2.25). Only participants for whom the full EEG and survey protocol was completed were included (total of one hour per participant). On average, 25.6 participants were recorded per device per week.

**Figure 2:**
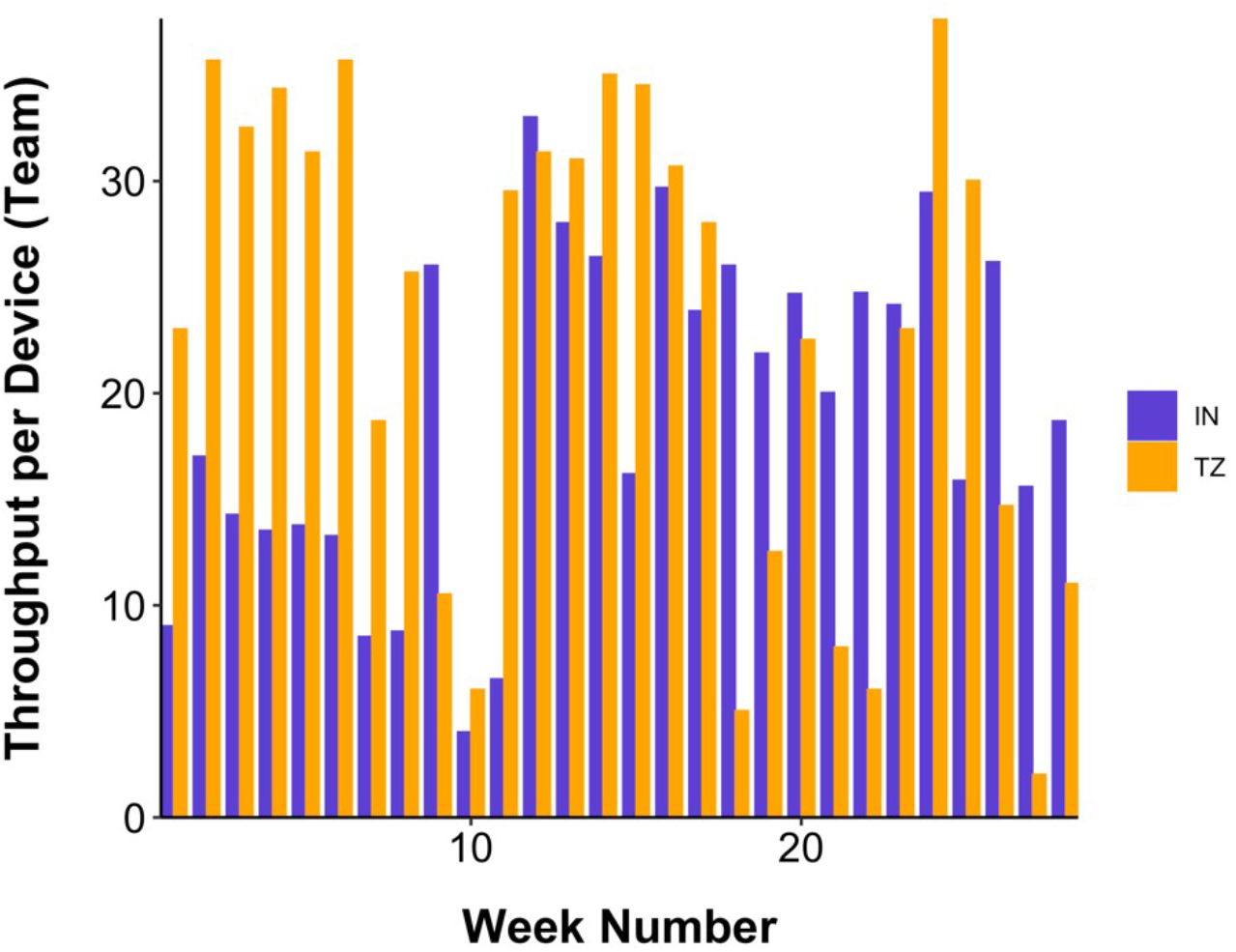
Weekly throughput for the India (IN) and Tanzania (TZ) pilot phase calculated as participants recorded per EEG device or research team.

### Percentage of bad channels

Figure 3 shows the mean percentage of bad channels based on PREP and FASTER for all data from Tanzania, India and each of the three benchmark datasets[BM1-BM3 (Singh et al., 2022; Wang et al., 2022; Miltiadous et al., 2023; Anjum et al., 2024; Xiang et al., 2024); Cumulative Distribution Functions, CDFs, shown in Supplementary Figure 1]. In the aggregate, the percentage of bad channels was higher in the field data compared to the benchmarks using the PREP method (though comparable in some cases) and generally lower in the field data using FASTER. Specifically, the percentage of bad channels in the Eyes Closed (EC) condition (Figure 3A) was slightly higher in the field samples compared to benchmarks using the PREP method (2.24±0.26% and 3.63±0.36% for Tanzania and India, respectively, and between 0.6% and 1.6% for BM1-BM3). On the other hand, the percentage of bad channels was higher but comparable between both field samples and benchmarks using FASTER (Tanzania: 4.88±0.26%; India: 6.42±0.29%; BM1: 6.04±0.73%; BM2: 6.28±0.59%; BM3: 6.77±0.70%).

**Figure 3:**
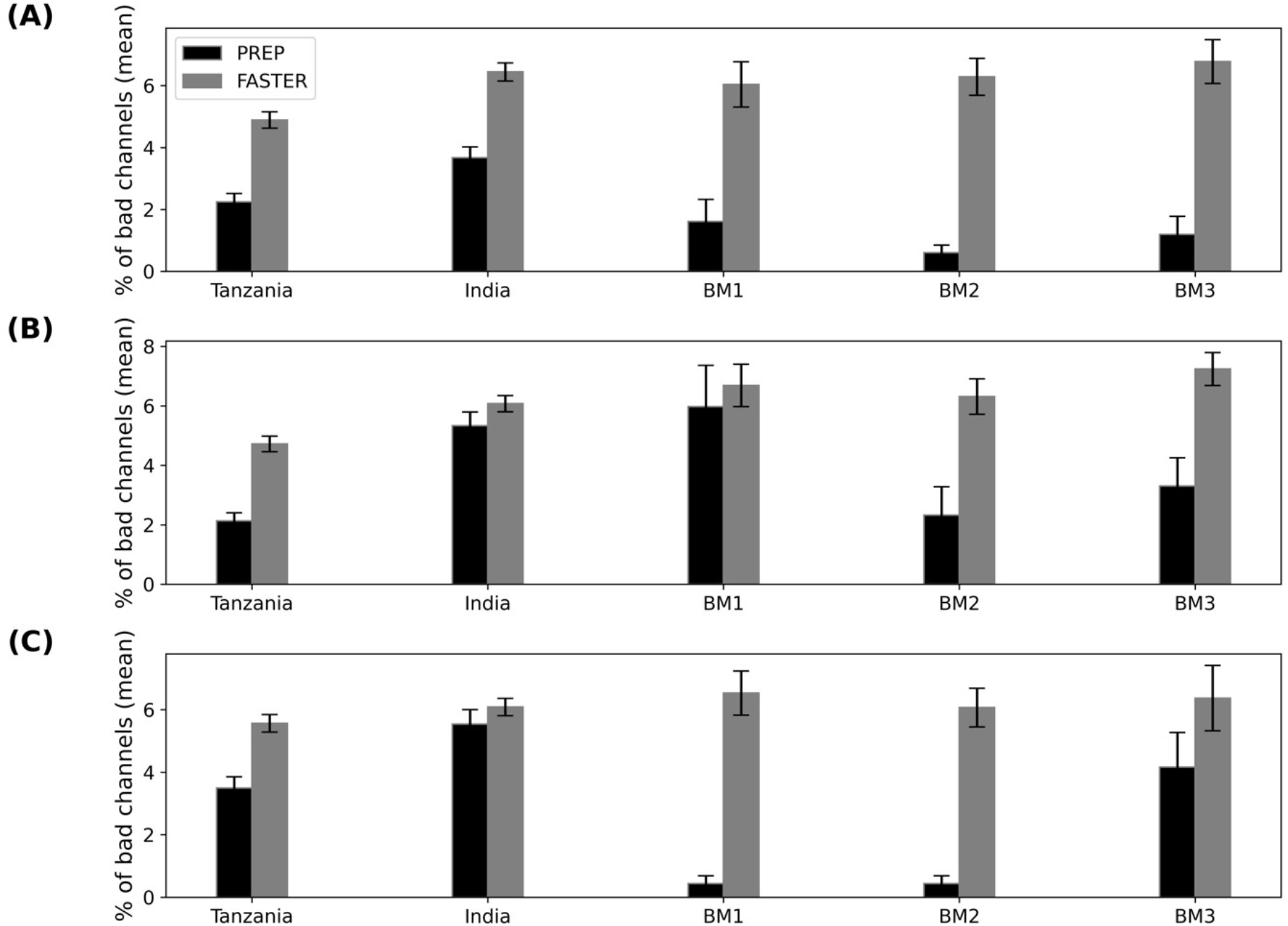
Average percentage of bad channels for Tanzania, India, and benchmark (BM1, BM2 and BM3) EEG recordings for Eyes closed (A), Eyes open (B) and Task Performance (C) conditions using PREP (black) and FASTER (grey) method. Error bars: standard error of the mean.

For the Eyes Open (EO) condition (Figure 3B), the percentage of bad channels was higher overall compared to EC for almost all datasets. Here the Tanzania data was similar to BM2 (Tanzania: 2.13±0.26%; BM2: 2.32±0.95%) and lower than BM1 (5.96±1.38%) and BM3 (3.30±0.94%), while India was higher (5.33±0.46%) but comparable to BM1 using PREP. Using the FASTER method, the Tanzania data had a lower percentage of bad channels (4.72±0.26%) compared to India and all the benchmark datasets (between 6 % and 8%), while the percentage of bad channels between the India data and the benchmarks were comparable (ranging between 6% and 7%).

In the case of the task performance condition (Figure 3C), while BM1 and BM2 had the lowest percentage of bad channels using the PREP method (0.43±0.24%), Tanzania and India datasets had comparable values to BM3, with Tanzania data having a lower percentage of bad channels (3.49±0.35%) compared to India (5.53±0.46%) and BM3 (4.16±1.1%). Using the FASTER method, we obtained comparable values across all datasets with the percentage of bad channels ranging between 5.5% and 6.5%.

### Percentage of bad epochs

Figure 4 shows the average percentage of bad epochs based on PREP and FASTER for all data from Tanzania, India and each of the three benchmark datasets (BM1-BM3; CDFs shown in Supplementary Figure 2). Here the field data had a lower percentage of bad epochs across almost all conditions using PREP, and a comparable percentage of bad epochs, compared to the benchmarks using FASTER.

**Figure 4:**
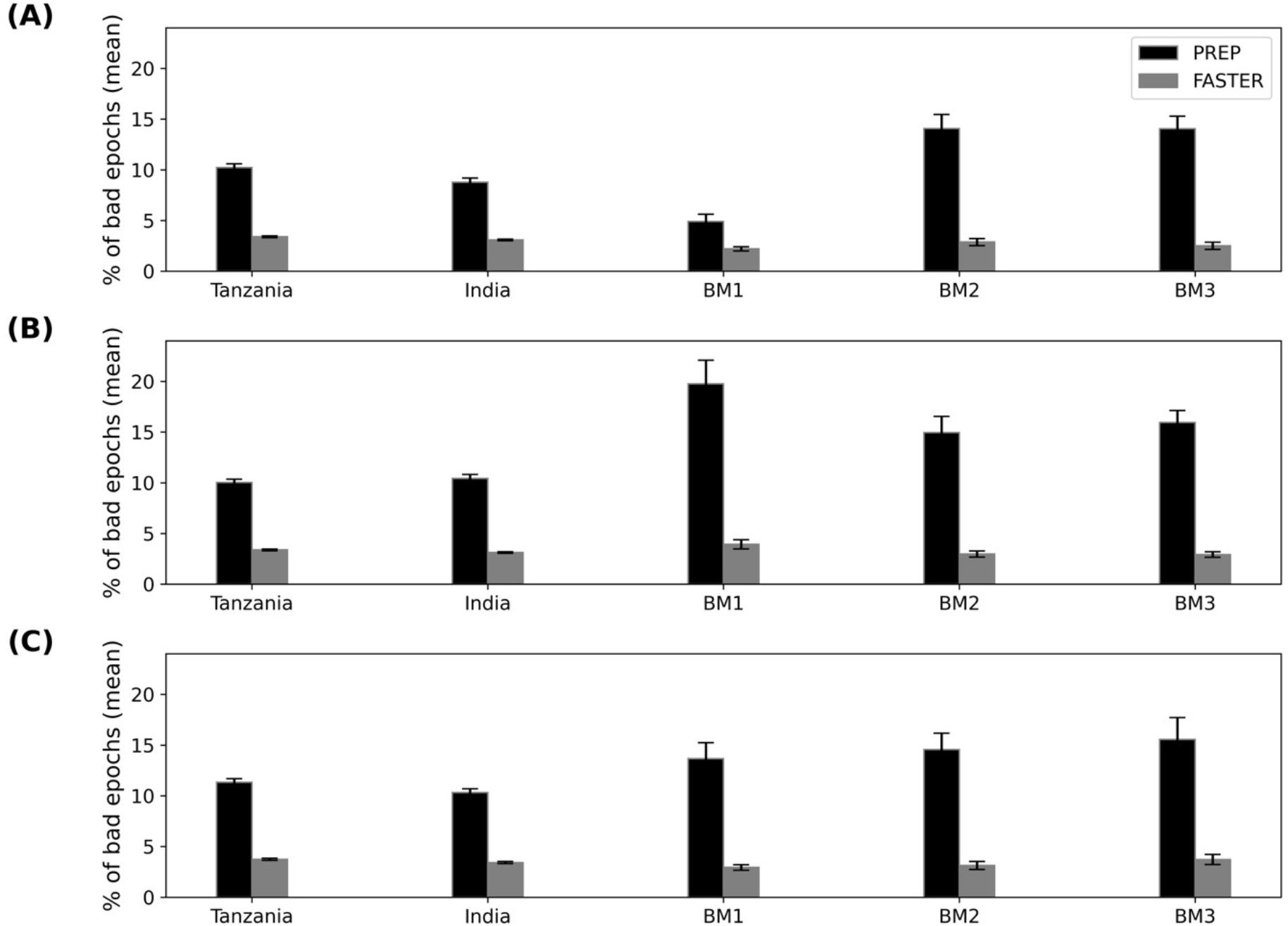
Average percentage of bad epochs for Tanzania, India, and benchmarks (BM1, BM2 and BM3). EEG recordings for Eyes closed (A), Eyes open (B) and Task Performance (C) conditions using PREP (black) and FASTER (grey) method. The error bars represent the standard error of the mean.

For the EC condition (Figure 4A) the percentage of bad epochs for Tanzania (10.21±0.37%) and India (8.77±0.40%) data was lower than BM2 (14.06±1.39%) and BM3 (14.06±1.22%) but higher compared to BM1 (4.90±0.71%) using PREP. Using the FASTER method, the results were comparable across all datasets with the percentage of bad epochs ranging between 2% and 3%.

For EO and task performance conditions (Figure 4B and 4C, respectively), using the FASTER method we again obtained comparable results for the percentage of bad epochs, with values ranging between 3% and 4%, while India and Tanzania datasets had a considerably lower percentage of bad epochs with the PREP method (EO-Tanzania: 10.03±0.32%; Task-Tanzania : 11.33±0.36%; EO-India: 10.42±0.39%; Task-India: 10.31±0.36%) compared to the benchmark datasets (ranging between 13% and 19%).

## Discussion

Here we have shown that it is possible to affordably collect high throughput, high quality EEG data across diverse field locations with training of an EEG naïve field research team coupled with robust systems and processes. This opens up a new frontier for research of diverse populations in diverse environments that is so crucially needed (Henrich et al., 2010; Dotson and Duarte, 2020). It also overcomes the practical and cost constraints associated with other types of neuroimaging infrastructure in low- and middle-income countries (Geethanath and Vaughan, 2019; Arnold et al., 2023).

### Data quality parameters

The percentage of bad channels in the field data was slightly higher than the benchmark data using PREP but comparable using FASTER. The percentage of bad channels was also higher in FASTER overall compared to PREP. This is because FASTER uses a standard amplitude Z-score of 3 as the threshold for detection, compared to a robust-Z-score threshold of 5 for PREP. This means that, while the field data had a comparable number of channels that met the >3 threshold, it had a larger number of channels that met the >5 threshold of the robust Z-score. We also note that the field EEG uses just 16 channels while the benchmark datasets use 64 channels. Thus, a single bad channel is equivalent to 6.25% in the field data but only 1.6% in the benchmark data.

In contrast, the percentage of bad epochs using PREP was comparable in the field data compared to two of the benchmark datasets but lower than one of them (BM1). The PREP method is traditionally used for bad channel detection and has been adapted to detect bad epochs, where if a robust Z-score of 5 for an epoch was exceeded in even one channel it is marked as a bad epoch. In FASTER the percentage of bad epochs was lower compared to PREP and comparable across the field and benchmark data. This is because it uses the average across all channels as the threshold for metrics such as amplitude range and variance. While channels may be eliminated due to artifacts, channels with bad epochs can still provide useful information. In fact, recent studies suggest that artifact removal can actually worsen results as it likely removes substantial regions of useful signal along with the artifacts (Delorme, 2023).

Finally, we also note several other factors of data quality beyond the EEG signal that have to be considered, such as correct capturing of channel names and other meta-data that is important for interpretation of the signal. While this is not shown here, these elements are also critically important aspects of the daily monitoring and feedback required to generate high quality data at scale.

### From lab to scale

Going from small to large scale without compromising data quality and doing so at reasonable cost is a challenge in many domains. While small lab studies are typically supervised by a PI along with students and post docs focused on the quality of their own study, large scale studies require a different paradigm. With a large number of people involved, the considerations for scale include standardization of methodologies, effective training methods and team structures as well as dashboards with daily analysis and feedback for rapid trouble shooting. The quality of data is thus more a reflection of the effectiveness of these processes over other factors such as researcher skill and device quality. In the absence of such processes, it is possible that issues may not be detected until much later, with data for many subjects having to be discarded. This will result in large costs due to wastage. With the throughput rates and data quality accomplished here, costs can be as low as $50/subject for a one-hour protocol that includes survey and EEG.

### The Sapien Center datasets

The large-scale data acquired here represents the pilot phase of an ongoing study that explores how changing human environments and the diversity of human experience differentially impacts brain physiology and functioning. In the first six months of this pilot, teams of 12 field researchers in two countries have already generated the largest database of general population EEG recordings in their countries or continents (N= 3,560 and N=4,373 for Tanzania and India respectively). The datasets include not just the EEG recordings described here but also extensive assessment of mental wellbeing or mind health along with a vast array of lifestyle, life experience and environmental factors.

## Data availability

We anticipate that this data will become dynamically available to the research community by the end of 2025 through our data platform Brainbase. This will include raw data as well as numerous standard and novel metrics computed from the EEG, along with survey elements. In the meantime, data is available on request.

## Acknowledgements

We thank members of the Sapien Labs team for their assistance with data management. We are grateful to all respondents for their participation in the project.

## Contributions

T.C.T. conceived the project. Sr.J-M.V. and S.Sw. supervised the projects in Tanzania and India, respectively. S.S.R. managed the field teams in India. S.P., J.V.A.K., P.T., and G.N.M.M. organized participant recruitment and location logistics in India. AK.P., K.G., S.K.A., A.A., V.D., P.R., O.S., Du.P., D.K., Pr.S., Pu.S., N.K., S.R., P.P. and S.Se. acquired field data in India. Sr.J-M.V., R.M. and A.S. organized participant recruitment and location logistics in Tanzania and managed the Tanzania field team. A.A.M., I.L.M., J.I.S., J.O.M., J.P.L., K.E.S., L.B.B., N.G.M., R.E.R., T.P.M., U.F.S., W.L.J. acquired field data in Tanzania. J.J.N. and Dh.P. assisted with EEG training of field researchers in India and Tanzania. N.P.S. performed the EEG benchmark analysis. Dh. P. conducted the throughput data analysis and designed the EEG dashboard. T.C.T., J.J.N. and N.P.S. wrote the paper with input from Sr.J-M. V., S.Sw., S.S.R. and Dh. P.

## Data Availability

The full dataset described in this paper is freely available for not-for profit research purposes. Access can be requested by emailing.

