## Supplementary Figure 1 for "EEG data quality in large scale field studies in India and Tanzania"

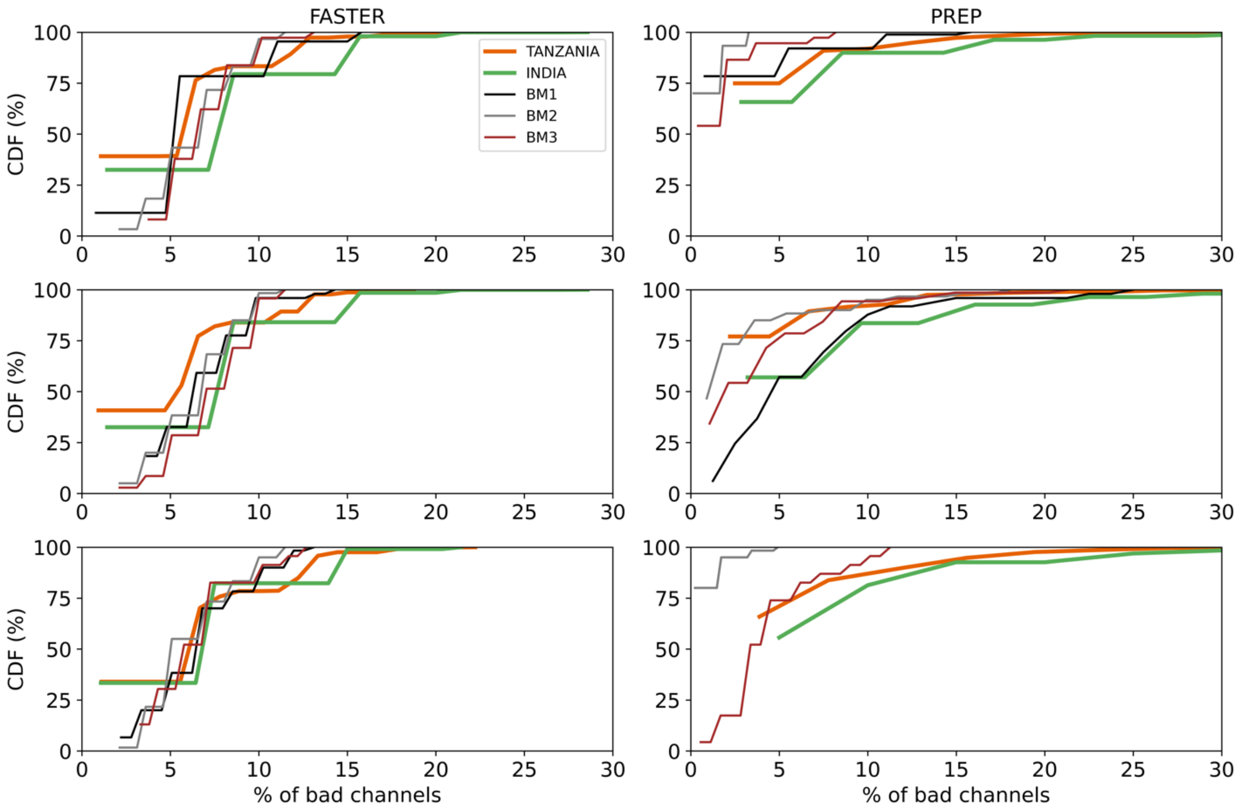


*Supplementary Figure 1: Cumulative distribution for the percentage of bad channels for FASTER (left) and PREP (right) for India, Tanzania and benchmark EEG datasets. Each row represents the EEG condition which include Eyes closed (top), Eyes open (middle) and task (bottom).*
